# Polar lipids are linked to nanoparticles in xylem sap of temperate angiosperm species

**DOI:** 10.1101/2021.10.22.465462

**Authors:** Xinyi Guan, H. Jochen Schenk, Mary R. Roth, Ruth Welti, Julia Werner, Lucian Kaack, Christophe L. Trabi, Steven Jansen

## Abstract

Xylem sap of angiosperm species has been found to include low concentrations of polar lipids and nanoparticles, including surfactant-coated nanobubbles. Although the nanoparticles have been suggested to consist of polar lipids, no attempt has been made to determine if nanoparticle and lipid concentrations are related. Here, we examined concentrations of nanoparticles and lipids in xylem sap and contamination control samples of six temperate angiosperm species with a NanoSight device and based on mass spectrometry. We found (1) that the concentration of nanoparticles and lipids were both diluted when an increasing amount of sap was extracted, (2) that their concentrations were significantly correlated in three species, (3) that their concentrations were affected by vessel anatomy, and (4) that concentrations of nanoparticles and lipids were very low in contamination-control samples. Moreover, there was little seasonal difference, no freezing-thawing effect on nanoparticles, and little seasonal variation in lipid composition. These findings indicate that lipids and nanoparticles are related to each other, and largely do not pass interconduit pit membranes. Further research is needed to examine the formation and stability of nanoparticles in xylem sap in relation to lipid composition, and the complicated interactions among the gas, liquid, and solid phases in xylem conduits.

## Introduction

One of the long-standing research questions in plant biology is how vascular plants transport water across xylem tissue over long distances (Hales, 1727). The question persists as to why plants are able to transport sap under negative pressure through dead xylem conduits without constant formation of large bubbles (embolism), which could reduce the transport efficiency considerably (Jansen and Schenk, 2015; Venturas *et al*., 2017). The assumption that high surface tension of xylem sap is required for a transpiration-driven process under negative pressure was recently challenged by the rediscovery of amphiphilic lipids in xylem sap (Scott *et al*., 1960, Esau 1965, Esau *et al*., 1966, Schenk *et al*., 2017, 2018, 2021). Moreover, the observation of nanoparticles and surfactant-coated nanobubbles in xylem sap raises questions about how plants are able to deal with a transport system that is subject to negative pressure but does not rely on pure and completely degassed liquid, as is typically required for artificial devices that operate under negative pressure (Wheeler and Stroock, 2008; Boatwright *et al*., 2015). Hence, more information on lipids and nanoparticles in xylem sap is needed to strengthen our understanding of their occurrence and functional significance.

Xylem sap is well known not to be pure water, but contains ions, sugars, proteins, lipids, etc. (Buhtz *et al*., 2004; Gollan *et al*., 1992; Gonorazky *et al*., 2012). A major challenge with chemical analyses of xylem sap is the extraction of sufficient sap material without contamination from cut living cells, including living fibres or parenchyma cells (Schurr 1988); this can be largely avoided by testing contamination control samples from the cut and cleaned surface of the xylem (Schenk *et al*., 2017, 2021). Very few lipidomic studies have been conducted on xylem apoplast fluid and/or xylem sap, but the findings so far show the presence of polar lipids, such as phospholipids and galactolipids, in xylem sap in all species tested to date (Gonorazky *et al*., 2012; Schenk *et al*., 2021). Schenk *et al*. (2021) found that the total polar lipid concentration varied from 0.18 to 0.63 nmol/mL across species, with monogalactosyldiacylglycerol (MGDG), digalactosyldiacylglycerol (DGDG), phosphatidic acid (PA), and phosphatidylcholine (PC) as the most common lipid types. These amphiphilic lipids are mostly attached to inner surfaces of conduits, including conduit walls and pit membranes, and are difficult to completely remove by sap extraction (Schenk *et al*., 2017, 2018, 2021). It has also been shown that xylem sap lipids are highly efficient surfactants, with dynamic changes of the surface tension depending on the local concentration of lipids per surface area (Yang *et al*., 2020).

Given that polar lipids may strongly affect the behaviour of gas-solid-sap interfaces at pit membranes, which are important safety valves between embolised and sap-filled conduits (Kaack *et al*., 2019, 2021), surfactant-coated nanobubbles in xylem sap are speculated to originate from the multiphase interactions at the pit membranes (Schenk *et al*., 2017; Yang *et al*., 2020). It is known that bubbles can be stable under negative pressure when their radius remains below a critical threshold, especially with the help of surfactants (Oertli, 1971; Atchley, 1989; Schenk *et al*., 2015). Evidence supporting this hypothesis includes the observation of nanoparticles in xylem sap based on a NanoSight device for five angiosperm species, and the visualisation of surfactant-coated nanobubbles in sap extracted from *Geijera parviflora* (Rutaceae) based on cryo-freeze fracture transmission electron microscopy (Schenk *et al*., 2017). While it is essential to test whether or not nanoparticles and nanobubbles occur in xylem sap of other species, it is also important to explore if these structures can be linked to xylem sap lipids. Since not all nanoparticles in xylem sap may represent nanobubbles, and because the NanoSight device is unable to distinguish nanoparticles from gas-core nanobubbles, we will use here the general term nanoparticles with respect to NanoSight experiments, and nanobubbles only if confirmed via electron microscopy.

In this study, we collected xylem sap from six temperate angiosperm species to conduct both nanoparticle and lipidomic analyses on the xylem sap extracted. In particular, we aimed to address the following three major questions.

(1) Is there any quantitative or qualitative correlation between lipids and nanoparticles in xylem sap? As gas-core nanoparticles coated with a mono-layer surfactant were observed in the xylem sap of *G. parviflora*, and the surfactant was speculated to be mainly amphiphilic lipids (Schenk *et al*., 2017), we expected a correlation between nanoparticle and lipid concentration. This relationship could also be affected by variation in lipid compositions, seasonal changes, and size distribution of nanoparticles.

(2) Does the lipid and nanoparticle concentration depend on the amount of xylem sap extracted? As pore sizes in pit membranes between conduits were reported to be well below 50 nm (Choat *et al*., 2003, 2004; Zhang *et al*., 2020), these would be too small for lipid micelles or nanoparticles whose diameter is typically > 150 nm to pass through (Schenk *et al*., 2017). As such, we predicted that most lipids in the extracted xylem sap would come from cut-open vessels at the cut end of a branch (Fig. 1; Schenk *et al*., 2021). We consequently hypothesize that there would be a dilution effect on lipid and nanoparticle concentration, with reduced concentrations for an increasing amount of xylem sap extracted. It is also possible that the concentrations of lipids and nanoparticles are affected by species-specific vessel anatomy, such as the cut-open vessel volume, and the total vessel perimeter at the cut xylem surface.

**Figure 1.**
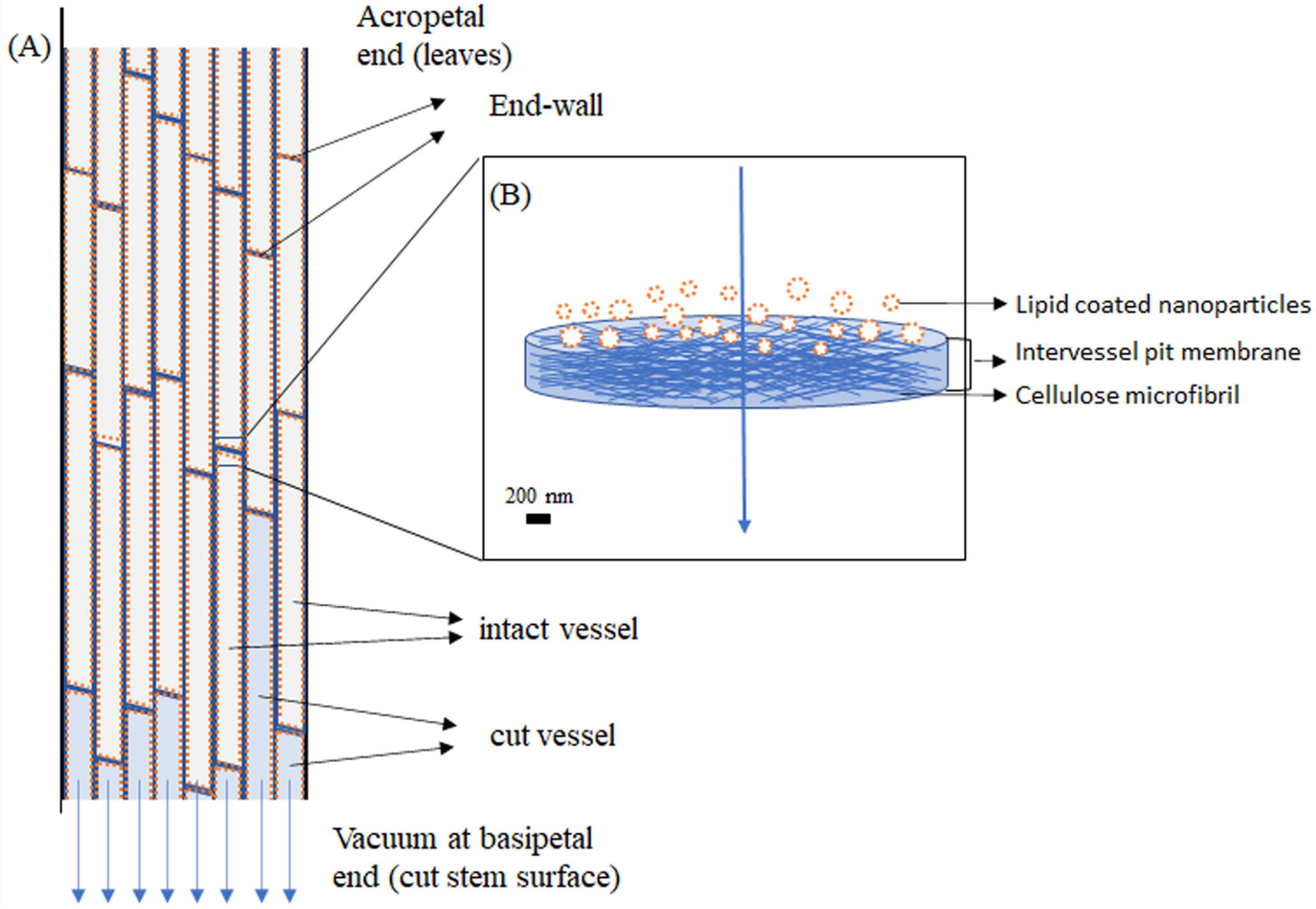
Diagrams of (A) xylem vessels in a stem subject to sap extraction by applying a vacuum at the acropetal end, and (B) the filtering capacity of an intervessel pit membrane in an end wall, which allows passage of sap, while lipid coated nanoparticles are largely prevented. Blue arrows indicate the direction of sap flow when applying a vacuum at the acropetal end of the stem. The orange dotted lines on the inner conduit walls (A) represent lipids, and also (B) coated nanoparticles. It is hypothesised that the lipids extracted in xylem sap originate only from the cut-open vessels (blue colour) at the acropetal end.

(3) Do lipids and nanoparticles occur as artefact-free compounds and objects in xylem sap of temperate angiosperm trees? Do seasonal changes have an impact on nanoparticles and lipids, and are nanoparticle dimensions and concentrations affected by a freeze-thaw cycle? We hypothesized that lipid and nanoparticle concentrations in xylem sap would be much higher than that in the contamination controls. Seasonal variation of nanoparticles and lipids is largely unknown, but clear seasonal variation in lipid composition has been found in *G. parviflora* (Schenk *et al*., 2021). Moreover, we expected that the mean nanoparticle size may vary among seasons and between fresh and defrosted sap samples.

## Materials and methods

### Plant material

Six deciduous angiosperm species, including *Betula pendula*, *Carpinus betulus*, *Corylus avellana*, *Fagus sylvatica*, *Liriodendron tulipifera*, and *Prunus avium* were studied. These represented common species at Ulm University (48°25’20.3” N, 9°57’20.2” E), Germany. For each species, four healthy and mature individuals were selected and labelled. Fresh branches were sampled and processed for measurements in August 2019, November 2019, February 2020, and July 2020.

### Xylem sap extraction

The extraction of stem xylem sap followed the methodology of Schenk *et al*. (2021), which investigated in detail the possibility of xylem sap contamination based on control samples, a lipid tracer experiment, and imaging. The experiments conducted by these authors showed that 95% of the xylem sap lipids did not originate from cut-open parenchyma cells at the basal stem surface.

Branch terminals with a length exceeding the maximum vessel length (approximately 1.5 to 2 meters) were collected during early mornings to ensure a fully hydrated status of the plants. The cut ends of branches were wrapped with moist tissue, and the branch terminals were covered with black plastic bags to prevent transpiration and transferred to the laboratory within half an hour. Immediately after arriving at the lab, a stem segment of ca. 10 cm was cut from the basipetal stem ends under deionized water to remove any potential embolism. Bark tissue was then removed from the cut end over a length of approximately 5 cm to expose the xylem and eliminate potential contamination from bark tissue. To further eliminate potential contamination, a clean and fresh razor blade was used to trim the cut surface, aiming to have cuts as clean as possible without obstructing the cut-open conduits. Cell debris and cytoplasmic remnants were removed using a high-pressure dental flosser (Ultra WP-100, Water Pik Inc., Colorado, USA) to thoroughly flush the exposed xylem cylinder with deionized water for about 90 seconds. A control sample was then collected for each branch to test for potential contamination prior to xylem sap extraction. The cut end of the xylem was submerged for 60 seconds into a watch glass containing 1 mL nano pure water, and the liquid was then transferred to a glass vial with a glass pipette. Contamination control samples were systematically collected during all experiments, except for the first sap extraction in August 2019.

Xylem sap was extracted by applying a vacuum to the basipetal end of a stem segment. The stem was inserted through a rubber flanged stopper, and the basipetal cut end was vertically placed into a glass tube embedded in a Buchner flask with ice. After checking the tightness of the whole system, we applied a vacuum to the flask. When the proximal end of the branch was under vacuum, all leaves were removed, and the branch was then shortened by removing acropetal ends. Therefore, we made ca. 5 cm successive cuts until sap was observed coming out from the basipetal end. Then, approximately 1 cm cuts were made successively until the desired sap volume was obtained. Xylem sap was transferred to glass vials and stored temporarily in a 4 °C refrigerator. The volume of the sap extracted was precisely recorded. The total amount of sap obtained from a single branch varied from 2 to 24 mL, which depended on the size of the branches, seasonality, and the xylem sap volume desired. Also, we paid special attention to working with clean glass vials to avoid contamination and to making the glass as hydrophilic as possible, which decreased the likelihood that amphiphilic lipids would stick to it during the lipid extraction process.

### Nanoparticle tracking analysis

Nanoparticles in xylem sap were investigated with a NanoSight LM10 microscope (Malvern Instruments Ltd., Malvern, UK). Nanoparticles show typical Brownian movements, which could be detected under a microscope when a laser beam with a wavelength of 638 nm was applied to the sample chamber. The concentration and size distribution of nanoparticles were calculated based on the number of nanoparticles per sap volume and their Brownian motion speed as detected with a NanoSight NTA 2.3 Analytical Software programme (Malvern Instruments Ltd., Malvern, UK).

Both fresh and frozen-defrosted xylem sap samples were included in the measurements. Fresh samples were stored at 4 °C immediately after extraction, and measurements were done within a week to avoid nanoparticle dissolution and microbial proliferation. Earlier work had shown that nanoparticles in xylem sap were stable over several days when kept refrigerated (Ushikubo *et al*., 2010). Because microorganisms show very different, non-Brownian movements, these could automatically be excluded from our analyses by the NanoSight software programme. Overall, we had very little or no contamination with microorganisms when our samples were stored in a refrigerator. Defrosted samples referred to samples immediately stored in a freezer (−18 □) after sap extraction, and defrosted at 4 °C one day before measurements were taken. We exanimated possible seasonal variation in nanoparticles by studying the frozen-defrosted xylem sap samples extracted in August 2019, and fresh xylem sap samples extracted in February 2020 and July 2020. To study the potential effect of freezing-thawing on nanobubble size and concentration, both fresh and defrosted xylem sap samples extracted in November 2019 were investigated and compared to each other.

### Mass spectrometry analysis

Mass spectrometry analysis of lipids was conducted on xylem sap and contamination control samples collected in August 2019 and February 2020, following the protocol of Schenk *et al*. (2021). One millilitre of fresh sap or control sample was freeze-dried using a SpeedVac concentrator unit (model Savant SPD121P, Thermo Scientific, Waltham, MA, USA) at 40 □. It took 2 to 3 h to get the samples completely dry. We then added 1 ml chloroform/methanol/water (5/5/1) solution to isolate lipids in the samples. The mixture was vortexed for 60 s to dissolve the solute fully, and centrifuged at 3,500 rpm for 8 min. The supernatant of the centrifuged sample was collected and transferred to a new 2 mL glass vial. This process was repeated twice, and the three supernatant solutions were merged. The supernatant solutions were put into the SpeedVac concentrator to freeze-dry, and the isolated lipids were then stored at −80 □. The lipid samples were sent with dry ice to the Kansas Lipidomics Research Center at Kansas State University for mass spectrometry analysis.

Direct-infusion electrospray ionization triple-quadrupole mass spectrometry was performed with a Waters Xevo TQS mass spectrometer (Waters, Milford, MA, USA). Lipid samples were dissolved in 1 mL chloroform, and the optimal amount of the lipid-chloroform solution (0.9-1 mL) was determined prior to the measurements. The samples were mixed with an internal standard mixture and 1.2 mL chloroform/methanol/300 mM ammonium acetate in water (300/665/35). The mixture was introduced to the mass spectrometer by continuous infusions with a 1 mL loop. Internal standards and mass spectral methods for the Waters Xevo instrument were as described by Schenk *et al*. (2021) with the addition of a neutral loss of 297.2 scan for 18:2-containing triacylglycerol (18:2-TAG) in positive mode as [M + NH_4_]^+^. The TAG internal standard was TAG (17:1/17:1/17:1). The cone voltage for 18:2-containing TAG analysis was 40 V and the collision energy was 6 V. Response factors to correct for differences in mass spectral responses of biological lipid molecular species compared to mass spectral responses of the internal standards were applied for lipid classes, except 18:2-containing TAG, which is reported as units of normalized mass spectral intensity, where an intensity of 1 unit is equal to the intensity of 1 nmol of internal standard.

### Cryo-freeze fracture transmission electron microscopy

Nanoparticles in xylem sap from *C. avellana* extracted with the method described above in June 2016, were visualised based on Papahadjopoulos-Sternberg (2010). Fresh sap was quenched using the sandwich technique and frozen with nitrogen-cooled propane, which avoided ice crystal formation and potential artefacts during the cryofixation process. The cryofixed sample was mounted on a standard Bal-Tec double replica holder, and inserted in a BAF 300 freeze-etching device (Balzers, Liechtenstein; Walther, 2003). The temperature in this device was raised to −120°C, and the vacuum was ca. 2 x 10^−7^ mbar. Fracturing was done by opening the double replica holder. The fractures were then coated with Pt for 30 s at an angle of ca. 30°. A second coating was done with carbon for 30 s at an angle of 0° (2kV per 60-70mA, 1×10^−5^ Torr). Unlike Schenk *et al*. (2017), the replicas were not cleaned by fuming concentrated HNO_3_ over the samples. The cleaning step may allow more detailed observations of the lipid coating, but was not required for the identification of nanobubbles. Two samples of xylem sap were prepared, and pure water was taken as a control sample. Observation was done with a JEOL JEM-1400 series 120kV Transmission Electron Microscope.

### Does the amount of extracted sap volume affect the lipid and nanoparticle concentration?

To test the potential impact of the amount of sap extracted on nanoparticle concentration, we extracted xylem sap following the steps described above. Instead of collecting sap from a branch in one tube, we collected the 1^st^, 2^nd^, 3^rd^, 4^th^, and 5^th^ mL of xylem sap separately in different glass tubes. Nanoparticle concentration was measured with a NanoSight LM10. One branch sample per species was used.

The mean nanobubble concentration of the multiple sap samples was calculated as follows,

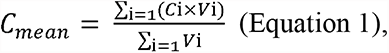

where *C* was the original nanobubble concentration, *V* was the volume of xylem sap extracted, *i* was the serial number of the sap sample investigated, and *C_mean_* was the average nanobubble concentration in the cumulative amount of sap extracted (Σ*Vi*). *C_mean_* was plotted against Σ*Vi* for each species to estimate the potential dilution effect.

The open vessel volume (mL), and the total vessel perimeter at the cut surface (cm) were calculated for each species:

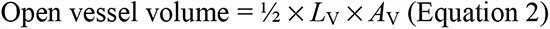

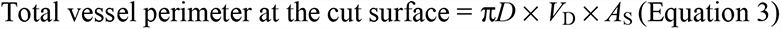

The average vessel length (*L*_V_, cm), average vessel diameter (*D*, μm), and vessel density (*V*_D,_ no mm^−2^) were retrieved from Guan *et al*. (submitted), which were based on the same tree individuals as studied here. Vessel area at the cut end of a branch segment (*A*_V,_ cm^2^) and stem xylem surface area (*A*_S_, cm^2^) were determined based on images of transverse sections of the stem segments collected in February 2020. Surface area was measured using the Fiji version of ImageJ (Schindelin *et al*., 2012).

Because the volume of sap extracted was found to impact the concentration of nanobubbles (Fig. 2A, Fig. S1), we normalised the nanoparticle concentration of each sap sample based on the species-specific logarithmic regressions to nanoparticle concentration per mL sap extracted.

**Figure 2.**
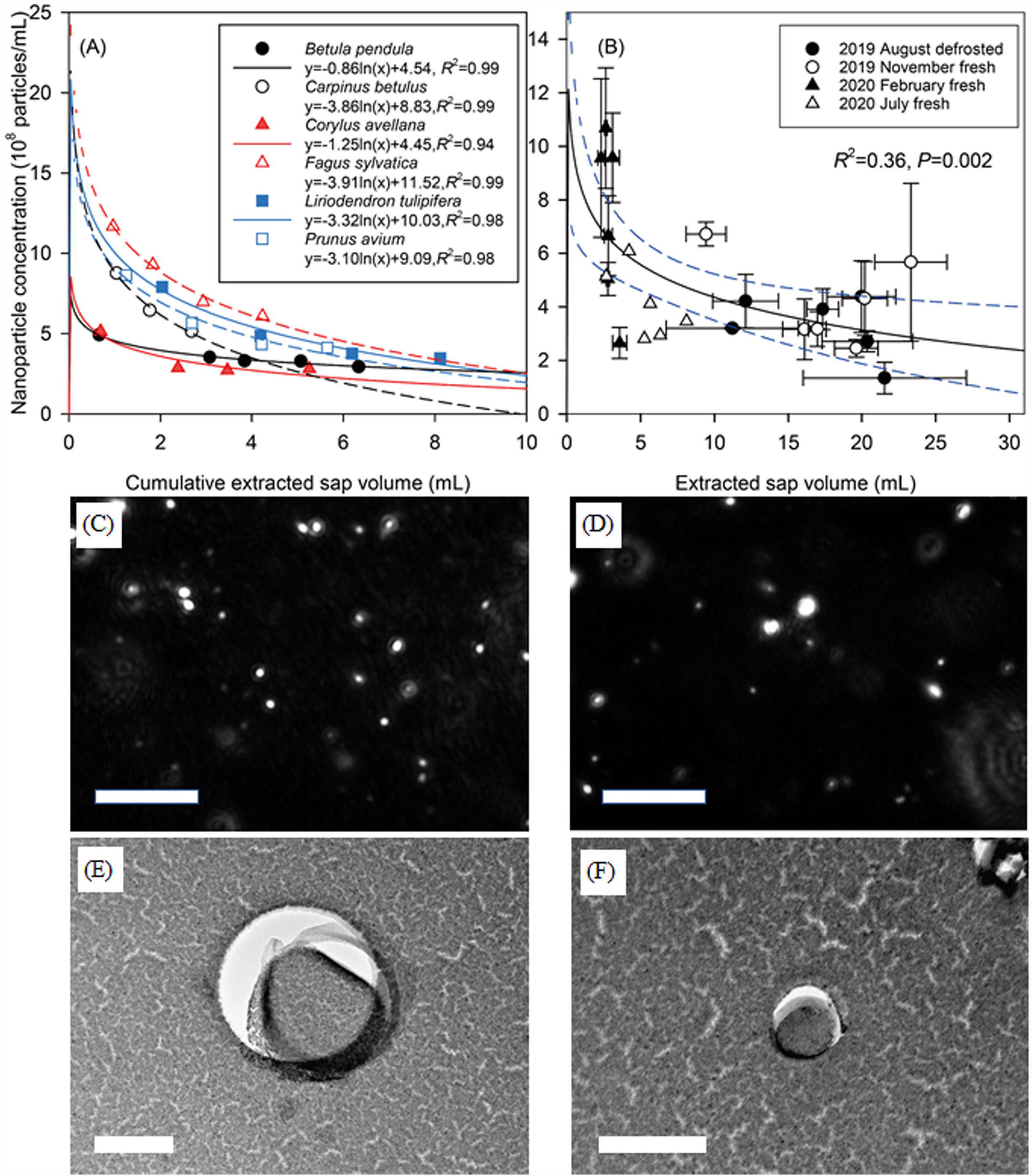
(A) Impact of the cumulative volume of extracted sap (mL) on the mean nanoparticle concentration (10^8^ particles per mL) based on consecutive sap extraction from a single branch per species in July 2020. Species are shown with different symbols and species-specific logarithmic regressions are calculated for further normalisation of sap samples per volume extracted. (B) Relationship between the volume of extracted sap (mL) and the nanoparticle concentration (10^8^ particles per mL) for sap samples extracted from different species and in different seasons. Data are presented as mean values ± standard deviation (n = 4 individuals per species) for samples collected in August 2019, November 2019 and February 2020; a single branch per species was collected in July 2020. The total sap volume and its corresponding mean nanoparticle concentrations in (A) were included in (B). Typical NanoSight images showing nanoparticles in the 1^st^ (C) and 4^th^ (D) sap sample during consecutive sap extraction from *Fagus sylvatica*. Freeze-fracture electron micrographs of surfactant-coated nanobubbles in xylem sap (E) and a control sample (F) of *Corylus avellana*. Nanobubbles are visible as white gas bubble cores (Pt/C-free areas) and dark, Pt/C decorated surfactant shells, which frequently shifted away from the bubble, and wrinkled during sample preparation. Scale bars: 20 μm for NanoSight images and 100 nm for ff-EM images.

### Data analysis

Species specific curves of nanoparticle concentration against sap volume extracted were plotted and fitted using SigmaPlot 14 (Systat Software Inc., Erkrath, Germany). After testing data for normal distribution and homogeneity of variance, normalised nanobubble concentration of sap collected in different seasons were compared using a one-way ANOVA for each species. Lipid compositions were obtained by converting lipid concentrations to percentages for each sample. Comparison of lipid compositions between xylem sap samples and contamination control samples, among sap samples of different species was made based on a principal component analysis (PCA). Changes in lipid compositions between summer and winter were examined using an independent sample t-test. Statistics were performed in SPSS 22 (IBM, Armonk, New York, USA) and R (version 4.0.3, R Core Team, 2016).

## Results

### Normalisation of nanoparticle concentrations in xylem sap and the dilution effect

The first millilitre of sap extracted from a branch showed the highest nanoparticle concentration for all species studied (Fig. S1). In contrast, lower concentrations of nanoparticles were found in the consecutive sap samples. Nanoparticle concentration in the latter samples was within the same range as the control samples (Fig. S1). We found a logarithmic decrease of mean nanoparticle concentration with increasing cumulative volume of sap extracted for each branch, and the species-specific regressions fitted very well to the measured data (0.94 ≤ *R*^2^ ≤ 0.99, Fig. 2A).

Moreover, a significant nonlinear correlation between nanoparticle concentration and extracted sap volume was also found in sap samples collected when analysing all species over the different seasons together (*R*^2^ = 0.36, *P* = 0.002, Fig. 2B). This also indicated that there was a dilution effect on the nanobubble concentration when a high amount of sap volume was extracted.

### Comparison of nanoparticle characteristics in xylem sap and contamination controls

To account for the dilution effect, nanoparticle concentration per mL sap extracted was normalised based on the species-specific regression equations shown in Fig. 2A. The normalised nanobubble concentration in fresh sap samples collected in November 2019 varied from 5.51 ± 2.43 10^8^ particles/mL for *B. pendula* to 16.03±2.32 10^8^ particles/mL for *F. sylvatica*. In contrast, nanoparticle concentration in the contamination control samples ranged from 0.60 ± 0.14 10^8^ particles/mL for *C. avellana* to 1.29 ± 0.61 10^8^ particles/mL for *L. tulipifera* (Table 1). Therefore, possible contamination from the cut surface of a stem could have caused 3% up to 35% of the nanoparticle amount detected in sap samples, with an average of 10% for all six species.

**Table 1.** Normalised nanobubble concentration (10^8^ particles per mL) of xylem sap samples and control samples. Values indicate means ± standard deviation (n = 4 individuals per species).

|  | 2019<br>August | 2019 November |  |  | 2020 February |  |
| --- | --- | --- | --- | --- | --- | --- |
|  | Defrosted<br>sample | Fresh<br>sample | Fresh<br>control | Defrosted<br>sample | Fresh<br>sample | Fresh<br>control |
| *B. pendula* | 6.31±1.98 | 5.51±2.43 | 1.18±0.31 | 6.35±2.22 | 5.92±1.20 | 0.57±0.41 |
| *C. betulus* | 14.90±1.47 | 13.90±0.83 | 0.68±0.20 | 15.06±1.01 | 14.44±4.68 | 0.54±0.14 |
| *C. avellana* | 6.15±1.15 | 9.50±0.81 | 0.60±0.14 | 6.73±1.19 | 10.60±6.05 | 0.72±0.19 |
| *F. sylvatica* | 16.01±3.00 | 16.03±2.32 | 1.06±0.28 | 14.45±1.41 | 10.66±3.30 | 0.94±0.17 |
| *L. tulipifera* | 11.46±0.94 | 12.57±1.29 | 1.29±0.61 | 11.13±1.11 | 6.82±0.70 | 0.90±0.26 |
| *P. avium* | 12.04±1.03 | 15.39±5.54 | 0.62±0.22 | 16.29±4.62 | 12.97±3.56 | 0.94±0.32 |

Similar results were found for the fresh sap and control samples collected in February 2020, where *B. pendula* showed the lowest normalised concentration of nanoparticles (5.92 ± 1.20 10^8^ particles/mL) and *C. betulus* showed the highest concentration (14.44 ± 4.68 10^8^ particles/mL, Table 1). Normalised concentrations of nanoparticles in the control samples varied from 0.54 ± 0.14 to 0.94 ± 0.32 10^8^ particles/mL, which accounted for 3 to 20% of the nanoparticle amount in the sap, with an average of 9% for the six species studied.

The mean diameter of nanoparticles in xylem sap ranged from 171 ± 18 nm for *C. avellana* to 232 ± 13 nm for *C. betulus* for the samples collected in November 2019, with an average of 202 ± 24 nm. The mean diameter of nanoparticles in the control samples showed a narrow range from 217 ± 5 nm (*C. betulus*) to 233 ± 10 nm (*F. sylvatica*), with an average of 225 ± 23 nm (Table S1). Similar results were found for the samples collected in February 2020, with nanoparticle diameter ranging from 189 ± 7 nm to 235 ± 13 nm for xylem sap of *P. avium* and *L. tulipifera*, respectively, and from 209 ± 12 nm to 243 ± 21 nm for control samples of *B. pendula* and *L. tulipifera*. The mean nanoparticle diameter was significantly larger in the control samples than sap samples (*F* = 0.047, *df* = 46, *P* = 0.002 for sap and control samples collected in August 2019; *F* = 0.161, *df* = 46, *P* = 0.004 for sap and control samples collected in February 2020).

### Seasonal variation and the effect of a freeze-thaw cycle on nanoparticle characteristics

Compared to fresh sap samples, the normalised concentration of nanoparticles in the frozen-defrosted samples collected in November 2019 did not differ in five out of the six species studied. However, a significant lower nanoparticle concentration was found in the defrosted sap of *C. avellana* as compared to the fresh sap (Fig. 3A, Table 1). A general increasing trend in nanoparticle diameter after a freeze-thaw treatment was found in all six species studied, but this difference was only statistically significant for *B. pendula* (Fig. 3B).

**Figure 3.**
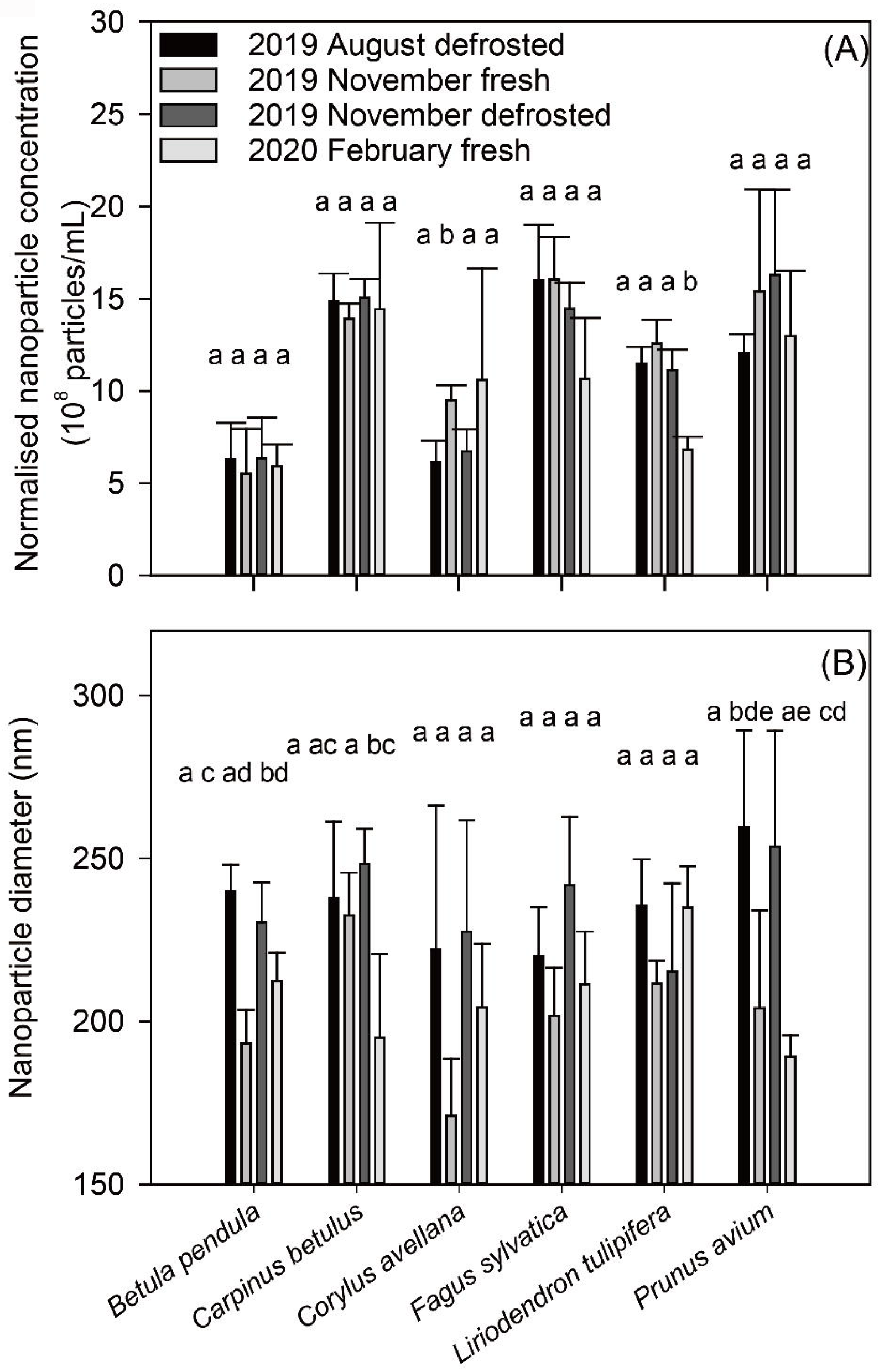
Variation in (A) normalised nanoparticle concentration (10^8^ particles per mL) and (B) nanoparticle diameter (nm) in xylem sap samples of different treatments and/or seasons. Data are presented as mean values ± standard deviation (n = 4 individuals), and different letters indicate a significant difference between samples (*P* < 0.05).

There was no significant difference in normalised nanoparticle concentration among sap samples collected over different seasons for *B. pendula*, *C. betulus*, *F sylvatica* and *P. avium*. Normalised nanoparticle concentration in fresh samples of *C. avellana* collected in November 2019 was significantly higher than the other samples collected from this species, while the fresh samples of *L. tulipifera* collected in February 2020 showed a significantly lower normalised nanoparticle concentration than other samples of *L. tulipifera* (Fig. 3A). No significant variation in nanoparticle diameter of xylem sap was found among all sample groups for *C. avellana*, *F. sylvatica* and *L. tulipifera*. The mean nanoparticle diameter was not significantly different between the frozen-defrosted samples collected in August 2019 and November 2019, and between the fresh sap samples collected in November 2019 and February 2020 for five out of the six species studied, indicating a lack of seasonal variation in nanoparticle diameter for these species. Significant seasonal difference in nanoparticle diameter was only found between the fresh sap samples collected in November 2019 and February 2020 for *B. pendula* (Fig. 3B).

### Lipid concentrations in xylem sap and contamination controls

For samples collected in February 2020, the total lipid concentration in xylem sap varied from 0.137 ± 0.036 nmol/mL (*P. avium*) to 0.376 ± 0.182 nmol/mL (*B. pendula*). The total lipid concentration was on average four times larger in sap samples than contamination control samples (Table S2). For samples collected in August 2019, the total lipid concentration varied from 0.071 ± 0.015 nmol/mL (*P. avium*) to 0.234 ± 0.071 nmol/mL (*B. pendula*, Table S2). Although the total concentration of lipids was significantly higher in February 2020 than in August 2019 (*F* = 5.100, *df* = 37.35, *P* < 0.001), a much smaller amount of sap was extracted in February 2020 (on average 2.88 ± 0.67 mL) than in August 2019 (on average 17.10 ± 5.45 mL, Table S1). Similar to nanoparticle concentrations, lipid concentrations in xylem sap samples collected in August were clearly diluted, and the lipid concentration in xylem sap was therefore only three times the lipid concentration in contamination controls (Table S2).

Based on the PCA analysis, there was a clear difference in the lipid composition between the sap samples and the contamination controls, with larger variation in lipid composition in control samples than in sap samples (Fig. S2, S3). Plant species were mainly grouped based on the loading scores of PCA analysis, that both summer and winter xylem sap samples showed similarities in lipid composition among different species (Fig. 5A, B).

**Figure 4.**
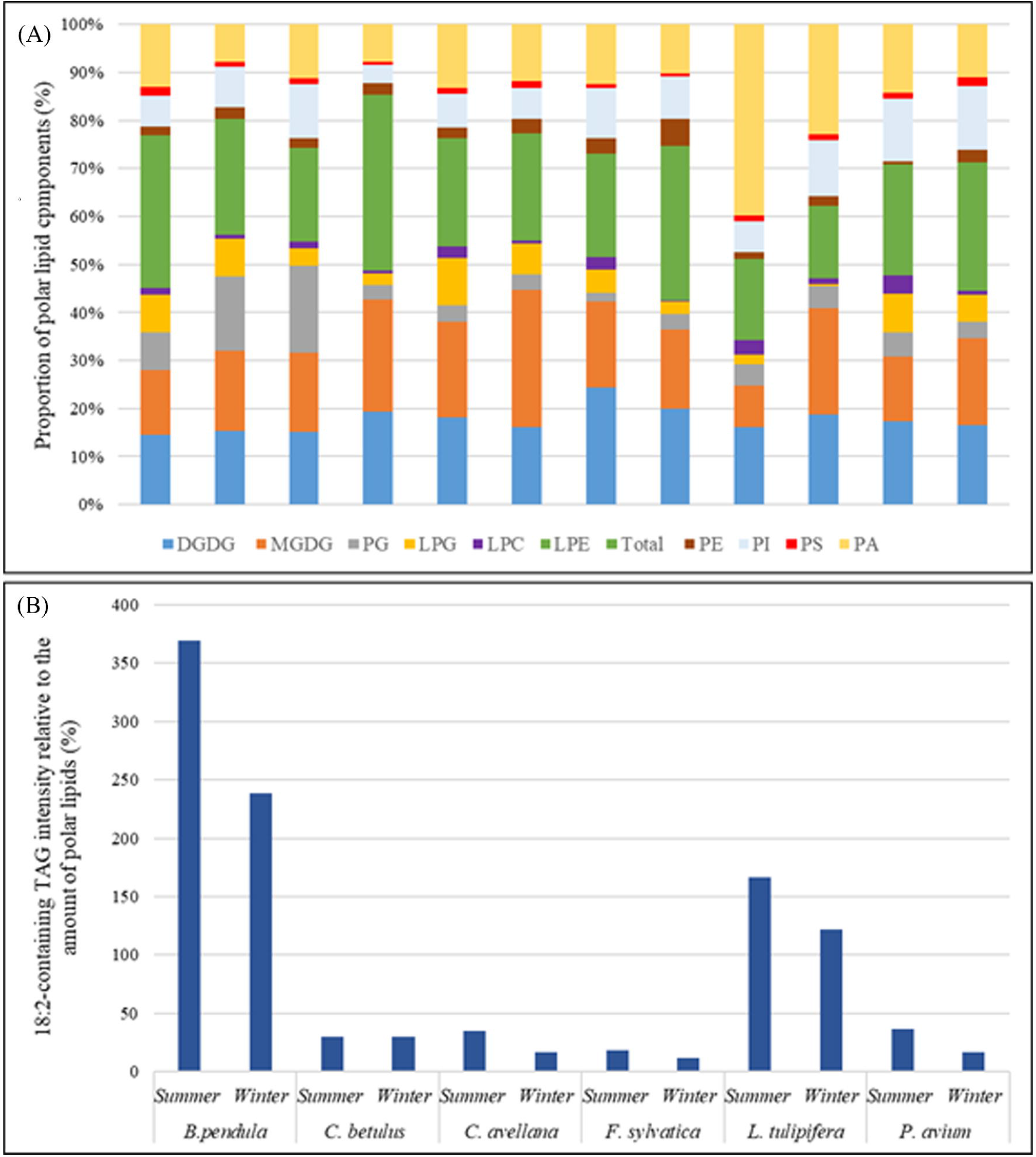
(A) Composition of polar lipids, and (B) 18:2-containing TAG in sap samples of six temperate angiosperm species extracted in August 2019 (summer) and February 2020 (winter), respectively. Galactolipids: DGDG = digalactosyldiacylglycerol, MGDG = monogalactosyldiacylglycerol. Phospholipids: LPG = lysophosphatidylglycerol, LPC = lysophosphatidylcholine; LPE = lysophosphatidylethanolamine, PA = phosphatidic acid, PC = phosphatidylcholine, PE = phosphatidylethanolamine, PG = phosphatidylglycerol, PI = phosphatidylinositol, PS = phosphatidylserine. Neutral lipid: 18:2-containing TAG = 18:2-containing triacylglycerol. The sequence of lipids in the bars is the same as the sequence of lipid compounds shown in the caption.

**Figure 5.**
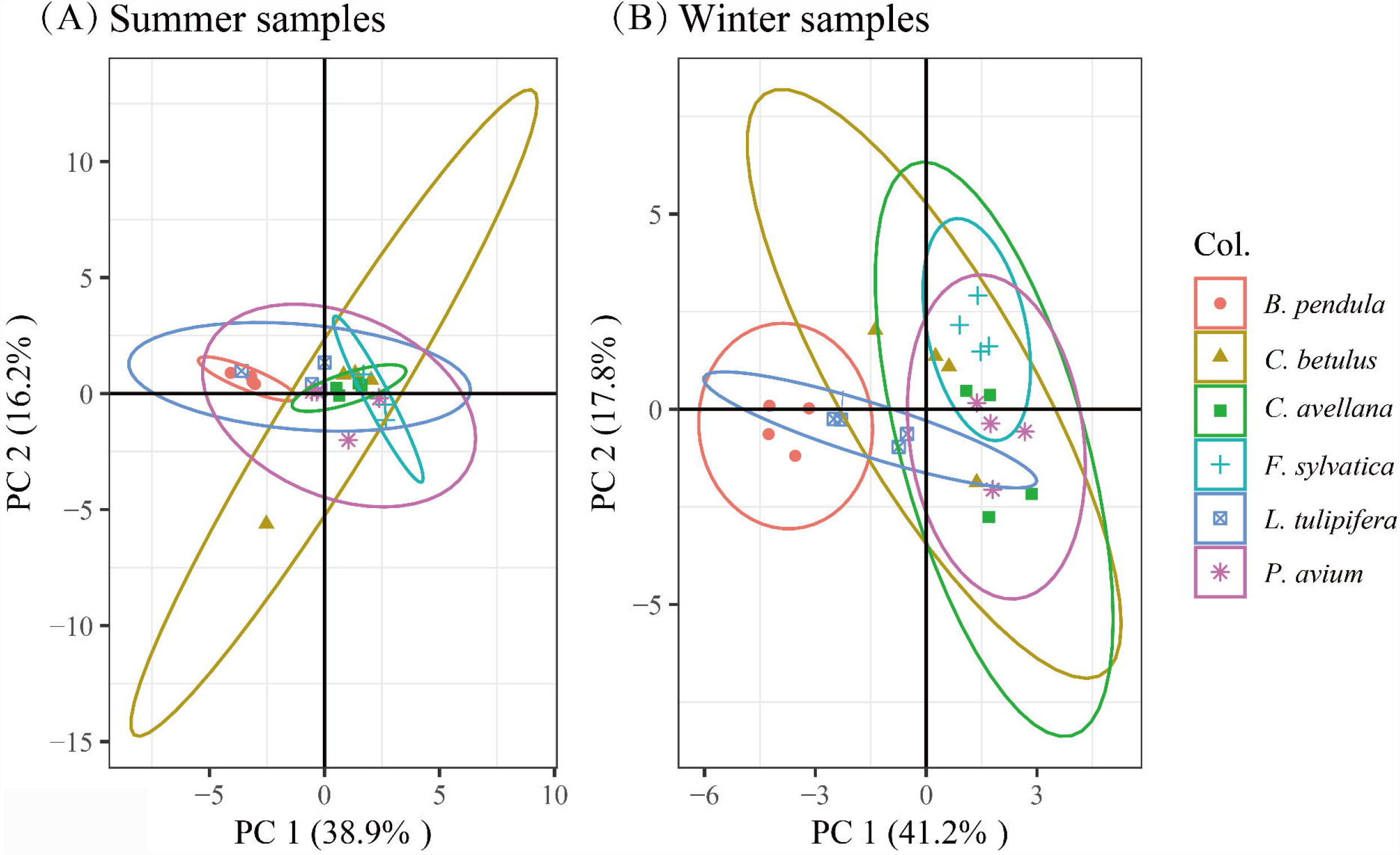
Scores plots of principal component analysis (PCA) based on lipid composition in percentage of (A) 2019 summer and (B) 2020 winter xylem sap of six angiosperm species. Dots were grouped by plant species, and the elliptical areas are 95% confidence regions.

### Lipid composition of xylem sap and seasonal variation

Polar lipids in xylem sap and contamination controls were mainly galactolipids (i.e., digalactosyldiacylglycerol, DGDG, and monogalactosyldiacylglycerol, MGDG) and phospholipids (Fig. 4A, Table S2). The concentration of phospholipids included mainly phosphatidylcholine (PC), phosphatidic acid (PA), and phosphatidylinositol (PI). In contrast, a small number of other phospholipids was detected, including phosphatidylethanolamine (PE), phosphatidylserine (PS), lysophosphatidylcholine (LPC), lysophosphatidylethanolamine (LPE) and lysophosphatidylglycerol (LPG) (Fig. 4A, Table 2).

The neutral 18:2-containing triacylglycerol (TAG) showed 2-4 times and up to 2 times of the normalized mass spectral intensity of polar lipids in the sap of *B. pendula* and *L. tulipifera*, respectively, while the 18:2-containing TAG intensity only represented ca. 20-40% of the total intensity of polar lipids in the other four species (Fig. 4B). A significant difference in the percentage of TAG out of the total normalized mass spectral intensity of all lipid was found between summer and winter xylem sap of *C. avellana* and *P. avium* (Table S3).

The xylem sap samples collected in August 2019 showed that the concentration of DGDG varied from 3.59 ±0.89% of the total lipids mass spectral intensity in *B. pendula* to 21.24 ± 2.48% in *F. sylvatica*, and MGDG varied from 3.61 ± 1.75% in *B. pendula* to 15.79 ± 3.03% in *F. sylvatica*. PA concentrations ranged from 2.93 ± 1.17 in *B. pendula* to 18.97 ± 12.59% in *L. tulipifera*, and PC varied from 6.67 ± 0.88% in *B. pendula* to 16.83 ± 3.35% in *C. avellana*, (Table S2). The lipid composition of sap collected in August 2019 was largely similar to the sap samples collected in February 2020, indicating a lack of seasonal change in most lipid compounds (Table S4, S5). Results of an independent sample t-test suggested no difference in lipid composition between summer and winter sap for *C. betulus* and *L. tulipifera* (Table S2). A significantly higher proportion of MGDG was found in the winter sap samples than in the summer ones of *P. avium*, and a higher proportion of PG was found in the winter sap samples than in the summer ones of *B. pendula* and *F. sylvatica*. The percentage of LPC in the winter samples was significantly lower than that in the summer samples for *C. avellana* and *F. sylvatica* (Table S2).

An increasing trend of total lipid concentration with increasing nanobubble concentration was found for several species and across seasons (Fig. S4). A significant and positive correlation between the original nanobubble concentration and lipid concentration was shown for three species, namely *C. avellana*, *L. tulipifera* and *P. avium* (Figure 6D-F). A similar trend was also observed in *B. pendula*, *C. betulus* and *F. sylvatica*, although this relationship was not statistically significant (Figure 6A-C).

**Figure 6.**
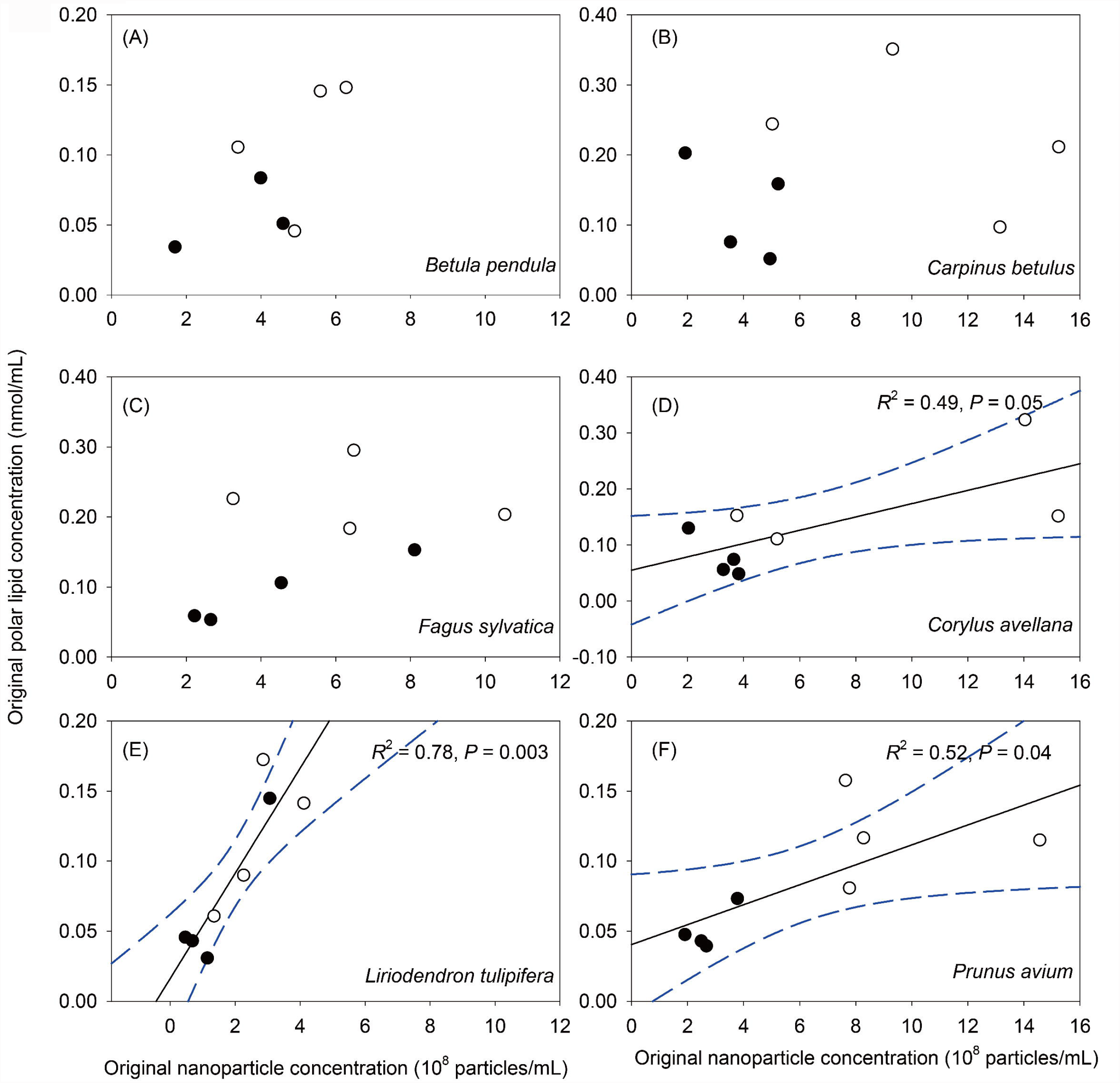
Relationship of original nanoparticle concentration (10^8^ particles/mL) and original polar lipid concentration (nmol/mL) for six angiosperm species. Each dot indicated one sample of xylem sap, which was used for both NanoSight and mass spectrometry measurements. The solid and open circles are xylem sap extracted in summer 2019 and winter 2020, respectively. The black solid and blue dashed lines indicated the linear regression and 95% confidence interval, respectively. The coefficient of determination and *P*-value was shown, when the correlation between the original nanobubble concentration and lipid concentration was significant.

### Linking lipid and nanoparticle concentrations with vessel anatomy

The nanoparticle and lipid concentrations were tested against vessel anatomical traits for the samples collected in February 2020 (Fig. 7). A significant, nonlinear relation was found between the polar lipid concentration and the open vessel volume, while the nanoparticle concentration was related to the total vessel perimeter at the cut stem surface.

**Figure 7.**
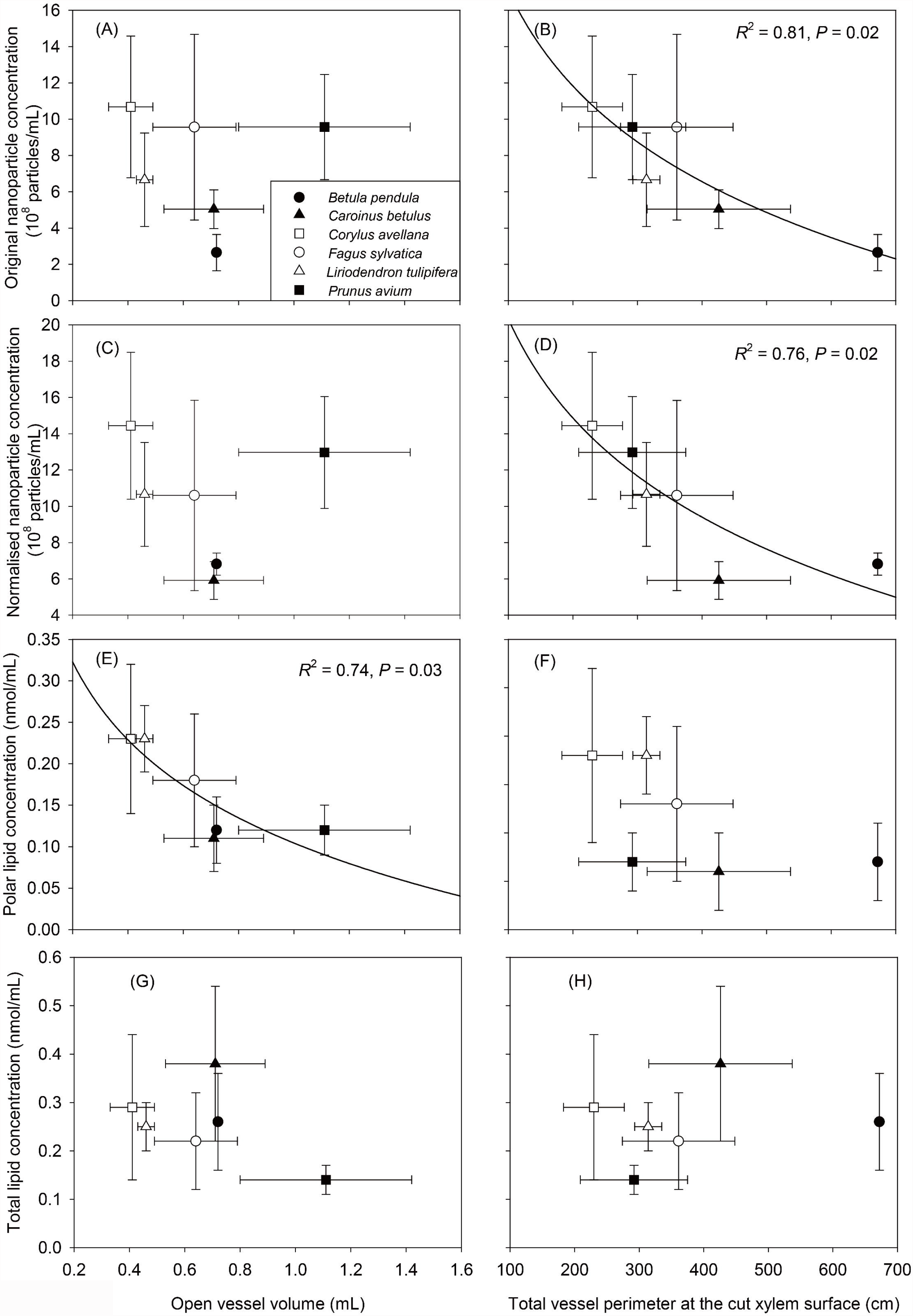
Relationship of vessel anatomy (open vessel volume, mL and total vessel perimeter at the cut xylem surface, cm) and nanoparticle/lipid concentration (original nanoparticle concentration, 10^8^ particles/mL; normalised nanoparticle concentration, 10^8^ particles/mL; polar lipid concentration, nmol/mL; total lipid concentration, nmol/mL). Each symbol represented the mean value of a species and its standard deviation. Solid lines showed a logarithmic regression.

### Visualisation of surfactant-coated nanobubbles in the sap of *Corylus avellana*

Freeze-fracture electron microscopy revealed coated nanobubbles in xylem sap of *C. avellana*. These were circular, but their shape was sometimes deformed, most likely due to sample preparation. The Pt and/or C-free areas appeared as white, inner gas volumes, while surfactant shells were dark due to Pt and C coating (Fig. 2E). The diameter of the nanobubbles varied from 43 nm to 243 nm, with an average size of 126 nm. The shells appeared as deflated, and frequently wrinkled structures, which were artificially removed from the bubble depression during sample preparation. Therefore, the shells overlapped the bubble partially in most cases, but some bubbles, especially the smaller ones, were still totally coated by complete shells. The nanobubbles had an isolated position, but some arrangements were observed. We also noticed a fairly dark, fuzzy layer on top of the surfactant shells, surrounding the nanobubble. The thickness of this layer was on average 20 nm. Additional xylem sap material may also stick as electron-dense structures to the nanobubbles.

Similar to the xylem sap samples, nanobubbles were found in the control samples (Fig. 2F). The diameter of nanobubble ranged from 37 nm to 183 nm, with a mean diameter of 63 nm, which was significantly smaller than nanobubbles in the xylem sap (Fig. 2E, F). Quantifying the number of nanobubbles based on freeze-fracture electron microscopy was not possible.

## Discussion

Although the surfactants of nanoparticles were speculated to consist mainly of amphiphilic lipids in previous studies (Schenk *et al*., 2017; 2021), a quantitative relation between polar lipids and nanoparticles has been demonstrated for the first time. Our results show that much higher concentrations of nanoparticles and lipids were found in extracted xylem sap of six temperate angiosperm species than in their contamination controls. Nanoparticles and lipids in xylem sap became diluted with increasing volume of xylem sap extracted, suggesting that most lipids extracted in the sap samples were not contaminations from living cells at the cut xylem surface, but largely came from cut-open vessels. Since a positive relationship between nanoparticle and polar lipid concentration was found for the six species studied, with a statistically significant relationship for three species, additional evidence is provided that polar lipids are the major components coating nanoparticles in xylem sap, with proteins possibly being another component. Moreover, the number of nanoparticles and lipid amount that could be extracted was also influenced by the open vessel volume and vessel surface, because lipids are mostly distributed on inner vessel walls.

While little seasonal variation was observed in nanoparticle concentration, size, and lipid compositions for most species studied, it remains unclear how nanobubbles in xylem sap behave and are affected by lipid composition and concentration in intact plants under a changing environment.

### Polar lipid and nanoparticle concentrations are related to each other

Significantly higher concentrations of nanoparticles lipids were found in xylem sap than in contamination controls, clearly suggesting that nanoparticles and lipids existing in xylem sap were collected by applying a vacuum to the basal end of a branch, and did not represent cell debris or contents from living cells (Schenk *et al*., 2017). The samples collected in November 2019 and February 2020 showed that the normalised nanoparticle concentration of xylem sap was on average 10 times higher than the contamination control samples (Table 1). The lipid concentration in xylem sap was also much higher than in the contamination controls (Table S2). Moreover, lipid compositions in the control samples varied considerably from branches of the same species (Fig S2, S3), suggesting quantitative and qualitative difference between lipids in the xylem sap and the contamination control samples.

Polar lipid concentrations generally increased with increasing values of original nanoparticle concentration (Fig. S4), and a positive and significant correlation of the concentration of polar lipids and nanoparticles was found for *C. avellana*, *L. tulipifera*, and *P. avium* (Fig. 6), suggesting a potential link between nanoparticles and polar lipids (Schenk *et al*., 2017). To date, the number of relevant studies on lipids and nanoparticles in xylem sap is limited. For the first time to our knowledge, we tested if the lipid and nanoparticle concentrations are correlated by using the same xylem sap samples. Interestingly, a mean nanoparticle concentration of 1.36 ± 0.48 10^8^ particles mL^−1^, and a polar lipid concentration of 0.18 ± 0.12 nmol mL^−1^ was found for xylem sap of *L. tulipifera* (Schenk *et al*., 2017; 2021). These values are very similar to the results in our study (Table 1, Table S2). Besides, high correlation coefficients between the concentration of nanoparticles and polar lipids were found consistently, and these were higher than the correlation between the concentration of nanoparticles and lipids, including neutral lipids (Figure S5). Although more species may need to be tested, these findings indicate that lipids and nanobubble characteristics could be species specific.

Triacylglycerol (TAG) containing the fatty acid 18:2 contributed more than half of the normalized mass spectra signal of lipids for *B. pendula* and *L. tulipifera* (Table S3). Unlike polar lipids, which are amphiphilic, TAGs are mainly hydrophobic and form lipid droplets in aqueous solutions. *Betula* has been characterized as a fat-storing genus, with tiny fat or lipid droplets detected with a light microscope in parenchyma cells (Sinnott, 1918; Harms and Sauter, 1992), as well as on inner vessel walls (Fig. S6, Westhoff *et al*., 2008). Increased TAG levels have been observed in plants under abiotic stresses and may serve as a reservoir for free fatty acids to avoid fatty acid-induced cell death in leaves generating free fatty acids (Xu and Shanklin, 2016). Lipid droplets also provide substrates and binding sites for lipid droplet-associated proteins, which synthesize bioactive compounds that can increase tolerance to biotic stresses (Lu *et al*., 2020). However, we are unaware of any research on neutral lipids in the plant apoplast, their functions, and potential changes under stress conditions.

Why are nanoparticle concentrations related to polar lipid concentrations? The presence of lipids in xylem sap, on inner vessel walls, and pit boarders has been cross-checked with the help of mass spectrometry, the fluorescent dye FM1-43 and OsO_4_ (Schenk *et al*., 2017; 2018; 2021). Nanoparticles in xylem sap has been observed in more than 10 species from various phylogenetic clades (Schenk *et al*., 2017; Schenk unpublished data and this study). The freeze-fracture electron micrographs of the nanoparticles also suggested that each nanoparticle owns a gas core coated with a surfactant shell (Fig. 2 E, F; Schenk *et al*., 2017). Therefore, it is plausible to speculate that nanoparticles in xylem sap are mainly coated nanobubbles. Xylem sap is never gas free, and the dissolution of gas in lipid phases may change due to temperature and pressure (Mercury *et al*., 2003). Embolism spreading from embolized vessels to sap-filled ones includes gas movement through intervessel pit membranes, either by mass flow or diffusion (Zimmermann, 1983; Sperry and Tyree, 1988; Kaack *et al*., 2019). Since pit membranes are confined, and lipid-coated, mesoporous media, nanobubbles would be coated with lipids during the penetration of gas-sap interfaces in pit membranes (Schenk *et al*., 2021). The coating of nanobubbles with lipids could reduce their surface tension, and consequently limit the size, which helps to prevent vessels from instant embolism (Schenk *et al*., 2015). The consistent relationship of nanoparticle and lipid concentration for all species was revealed quantitatively in our study (Fig. 6, Fig. S4), which adds new evidence that the surfactant coat of nanobubbles in xylem sap are mainly polar lipids.

The finding that polar lipid concentrations in xylem sap did not correlate tightly with nanoparticle concentrations in all species (Fig. 6) suggests that the nanoparticles may contain other components, most likely proteins (Buhtz *et al*., 2004), which are abundant in xylem sap residue (Schenk *et al*., 2017). Many proteins bind to lipids, especially in membranes, and non-specific lipid transfer proteins have been found in xylem sap of many plant species (Buhtz *et al*., 2004; Djordjevic *et al*., 2007; Krasikov *et al*., 2011; Ligat *et al*., 2011). As in lung surfactants (Parra and Pérez-Gil, 2015), proteins could play important roles in the behaviour of lipid surfactant layers in plant xylem. An additional explanation could be that lipids are difficult to extracted from inner conduit walls of cut-open vessels (Schenk *et al*., 2021).

### The concentration of nanoparticles and lipids is affected by the extracted sap volume and vessel anatomy

The mean diameter of nanoparticles ranged from 170 to 250 nm based on the NanoSight measurements, which is much larger than the pore sizes of pit membranes below 50 nm (Zhang *et al*., 2020). Therefore, nanoparticles are unlikely to pass through pit membranes, and the nanoparticles collected would mainly originate from the cut-open vessels at the base of our stem samples (Fig. 1). Since the volume of cut-open vessels ranged from 0.4 to 1.1 mL for the species studied, the highest nanoparticle concentration was always measured in the first mL of xylem sap extracted during the consecutive sap extraction from the same branch (Fig. 7, Fig. S1).

The mean nanoparticle concentration decreased rapidly with increasing sap volume, especially within the first several millilitres (Fig. 2A, B). Thus, significantly higher original values of nanoparticle concentration were detected when on average 3 mL of sap was extracted in February 2020 than the ca. 20 mL of sap extracted in August and November 2019 (Table S1). Similar results were also found for the lipidomic analyses based on the same samples collected in August 2019 and February 2020 (Table 2). Instead of seasonal changes in nanoparticle concentration, we identified that variation in the original nanoparticle concentration was mainly due to the sap volume extracted. Indeed, there was no longer any difference in nanoparticle concentration after data normalisation (Fig. 3A).

Lipids are insoluble in xylem sap and are found to mainly distribute on inner-conduit walls and pit borders (Schenk *et al*., 2018). A significant and negative correlation was found between nanoparticle concentration and total vessel perimeter at the cut xylem surface, and between polar lipid concentration and cut-open vessel volume in xylem sap collected in February 2020 (Fig. 7). A similar relation was reported between the polar lipid concentration and the total vessel perimeter at the cut xylem surface for another seven species (Schenk *et al*., 2021). Our results that fewer lipids could be extracted with larger vessel volume or surface agreed with the suggestion of Schenk *et al*. (2021) that polar lipids are difficult to be completely removed from vessel lumen surfaces. Since polar lipids are main components of the coating of nanoparticles, our observation that considerable nanoparticles were detected in the second and third mL of xylem sap extracted (Fig. S1), provides additional evidence to this hypothesis.

Further research aiming to quantify lipids based on mass spectrometry imaging of vessels before and after xylem sap extraction could help to understand lipid distribution in vessels and how tightly they attach to vessel walls (Ellis *et al*., 2013).

### How do plants cope with changing seasons with respect to nanoparticles and lipids?

Although the normalised concentration of nanoparticles showed little variation in the sap samples collected across different seasons, some variations of the nanobubble diameter were found among seasons and between fresh and defrosted sap samples (Fig. 3). To test if temperature would affect the stability of nanoparticles in xylem sap, a frozen-defrosted treatment was applied to the sap samples collected in November 2019, which simulated winter frost. Generally, larger nanoparticles were observed in the defrosted sap compared to the same fresh sap samples, but this difference was only statistically significant for *B. pendula* (Fig. 3B). Dissolved gas is released from sap and forms bubbles when sap freezes. The stability of these bubbles and their ability to be redissolved into the liquid phase during the thawing process could influence embolism formation (Sperry and Sullivan 1992). Lipid-coated nanobubbles could stabilise the existence of bubbles in the xylem sap, and the area-dependent surface tension also increases with increasing bubble diameter, which allows bubbles larger than the critical bubble size to be stabilised under certain conditions without requiring more lipids (Schenk *et al*., 2017; Yang *et al*.,2020). Although plants live in a much more complex and constantly changing environment, it is impossible for us to know if (nano)bubbles exist in intact plants, and how exactly they behave under negative pressure. More studies on nanobubble characteristics undergoing a freeze-thaw cycle may help to understand the potential role of lipids in preventing freeze-induced embolism.

An overall similarity in lipid composition was found between sap samples of August 2019 and February 2020. The mass spectra intensity of 18:2 containing TAG in winter sap differed significantly from the summer sap for *C. avellana* and *P. avium* (Table S3), but not for the other four species. This is in line with a previous study showing little seasonal variation of TAG concentration in the wood of *B. pendula* (Piispanen and Saranpää, 2004). Regarding polar lipids, MGDG, DGDG, and PA are the major components that consist of more than half of the total polar lipid concentrations (Fig. 4A). A significant change in the proportion of MGDG between summer and winter sap samples was only found in *P. avium*, but no significant change was found in the proportion of DGDG for all species studied (Table S3).

In a previous study (Schenk *et al*., 2021), the lipid composition in xylem sap was found to be very similar to the composition of lipids in the entire xylem, suggesting that there is no selective transport of lipids into vessels from surrounding cells. Instead, it seems likely that lipids in vessels originate from the living vessel content (Scott *et al*., 1960; Esau 1965; Esau *et al*., 1966; Schenk *et al*. 2021), and therefore do not change much in composition over time once the vessel becomes functional. Seasonal change in lipid composition has yet only been tested for *G. parviflora*, where a significant difference in the concentrations of DGDG and MGDG was found between xylem sap sampled in spring and summer. This could be due to sampling at different stages of xylem development (Schenk *et al*., 2021). Moreover, seasonal changes in the electron density of pit membranes have been observed (Schmid and Machado, 1968; Sorek *et al*., 2021), suggesting a potential lipid accumulation in membranes over time. Other experimental techniques, such as matrix-assisted laser desorption/ionisation MS imaging (MALDI), may help to visualise lipid distribution in xylem directly (Saito *et al*., 2012), and determine if there are any changes related to season or ageing of vessels.

## Abbreviations

DGDG: digalactosyldiacylglycerol
MGDG: monogalactosyldiacylglycerol
LPG: lysophosphatidylglycerol
LPC: lysophosphatidylcholine
LPE: lysophosphatidylethanolamine, PA, phosphatidic acid
PC: phosphatidylcholine
PE: phosphatidylethanolamine
PG: phosphatidylglycerol
PI: phosphatidylinositol
PS: phosphatidylserine
TAG: triacylglycerol

## Acknowledgments

We thank Andrea Huppenberger and Clara García Sánchez for lab assistance, various colleagues from the botanical garden for assistance with sampling plant material, and the work group Neurochemistry and Neurodegeneration at Ulm University for technical support with NanoSight observations. Paul Walter from the Electron Microscopy facility at Ulm University is acknowledged for assistance with cryo-freeze fracture transmission electron microscopy. This study was funded by the Deutsche Forschungsgemeinschaft (DFG, German Research Foundation, project nr. 410768178). H.J.S., R.W. and S.J. acknowledge financial support from the National Science Foundation (IOS-1754850).

## Author contributions

X.G. conducted experiments, performed formal analysis, curated all data, and wrote the original draft; R.W. and M.R. conducted lipidomic experiments and data analysis. H.J.S. and S.J. conceived and supervised the project. J.W., L.K. and C.T. performed part of the experiments. R.W., M.R., H.J.S. and S.J. edited and finalised the manuscript. All authors read and approved the final manuscript.

## Data availability

The data supporting the findings of this study are all included in figures, tables and supplementary material files

